# Kokiri: Random-Forest-Based Comparison and Characterization of Cohorts

**DOI:** 10.1101/2022.08.16.503622

**Authors:** Klaus Eckelt, Patrick Adelberger, Markus J. Bauer, Thomas Zichner, Marc Streit

**Affiliations:** Johannes Kepler University Linz; Boehringer Ingelheim RCV GmbH & Co KG

**Keywords:** Human-centered computing, Visual analytics, Applied computing, Genomics

## Abstract

We propose an interactive visual analytics approach to characterizing and comparing patient subgroups (i.e., cohorts). Despite having the same disease and similar demographic characteristics, patients respond differently to therapy. One reason for this is the vast number of variables in the genome that influence a patient’s outcome. Nevertheless, most existing tools do not offer effective means of identifying the attributes that differ most, or look at them in isolation and thus ignore combinatorial effects. To fill this gap, we present Kokiri, a visual analytics approach that aims to separate cohorts based on user-selected data, ranks attributes by their importance in distinguishing between cohorts, and visualizes cohort overlaps and separability. With our approach, users can additionally characterize the homogeneity and outliers of a cohort. To demonstrate the applicability of our approach, we integrated Kokiri into the Coral cohort analysis tool to compare and characterize lung cancer patient cohorts.

## 1 Introduction

With advances in cancer therapy, it is becoming increasingly important to study information beyond type and stage of a tumor and patient demographics. Newer drugs directly target specific cancer-causing gene mutations, enabling treatment that is specifically tailored to the individual patient [19]. However, studies have also shown that the gene mutations involved differ not only across cancer types, but also across race and sex [25,32]. This results in variations in therapeutic outcome across patient cohorts. Uncovering the reasons for these differences is a challenge because of the complexity of the problem and the amount of data to be considered. Thousands of genes, several hundred of which have been identified as tumor-relevant [34], may play a role. Knowledge on how patient cohorts differ and what the most important differences are is key to further improving therapy.

Large-scale projects, such as the AACR Project GENIE [1], provide clinical and genomic data about thousands of patients from which clinically relevant subtypes can be derived. Reasoning about such large amounts of data is becoming increasingly complex, and analysts require powerful tools to gain insights [12]. Visual analytics tools enable domain experts to visually explore and analyze cancer cohorts in such large datasets [2,8,13,35]. Comparison of cohorts, however, is often limited to single attributes or to multiple attributes that are being considered individually. Combinatorial effects of multiple attributes are thus lost.

To address these challenges, we propose an interactive visual analytics approach called Kokiri, which allows users to compare cohorts by their high dimensional data with the goals of (i) uncovering driving differences between them and (ii) characterizing the homogeneity of individual cohorts. We achieve this by training a random forest model to classify the cohorts based on their high-dimensional data. We report the most important attributes that differentiate between the cohorts and give an overview of the model’s classification performance. Users can iteratively refine the classification by limiting the data to a subset of interest, for instance, by excluding genes that are already known to differ between cohorts, to focus on the remaining data and gain further insights. Additionally, the homogeneity of individual cohorts can be assessed based on the classifier’s confidence, and hard-to-classify items can be identified. To demonstrate the utility of our approach, we describe a use case where we compare lung cancer cohorts to verify findings from literature and gain deeper insights into the data from the AACR Project GENIE [1].

## 2 Related Work

Comparing cohorts is a fundamental task in cancer research as well as many other domains that can be supported effectively by visual analytics tools. While *comparison* of cohorts looks for differences between two or multiple cohorts, *characterization* of a cohort looks for potential differences within a cohort.

Coral [2] is a cohort analysis tool specifically designed for creating and characterizing cohorts. However, visual comparison of cohorts is limited to one or two attributes that must be selected manually by the user. Coral integrates TourDino [10] for pairwise statistical comparison of cohorts by a user-defined set of attributes. In previous work, we have analyzed cohort differences in low-dimensional embeddings to provide an overview of differences in high-dimensional space [11]. Summary visualizations explain the data of individual cohorts, and difference visualizations highlight differences in attribute distributions between cohorts. Both visualizations are ranked such that the most and least varying attributes can be seen at a glance. Similarly, Duet [21], shows the most similar and different attributes for two selected cohorts and uses textual explanations alongside histograms as summaries of these similarities/differences. Differences in data distributions were also considered by Gotz et al. [14]. Their approach seeks to avoid the introduction of selection bias when creating cohorts. This bias is determined by calculating the Hellinger distance between pairs of cohorts. Building on this, Borland et al. [4] presented a set of visualizations to compare a user-defined focus cohort with a baseline cohort. In contrast to the approaches above, Somarakis et al. [33] described a system for analyzing single-cell omics data where cohorts can be compared based on the abundance of different cell types and combinations thereof. The system ranks attributes by their cohort-differentiating ability and also supports visual detection of outliers within each cohort. Except for the work of Somarakis et al. [33], all of the approaches above consider attributes only individually when comparing cohorts. In doing so, they ignore the vast combinatorial space of high-dimensional data, from which further differences can be gleaned. Furthermore, only Coral [2] allows comparison of more than two cohorts.

In machine learning, classification models categorize items into classes based on their data. Classification models take advantage of high-dimensional data to be able to distinguish between classes, and provide an alternative to traditional pairwise statistics as used in the approaches above. In our case, the cohorts are the classes that are to be distinguished. Visual analytics tools can support domain experts in constructing and analyzing classification models. Endert et al. [12] reviewed the state-of-the-art approaches to integrating machine learning methods into visual analytics workflows and also discussed how classification algorithms and people can work together. In BaobabView [38], domain experts can interactively create a decision tree to integrate their domain knowledge into the classification model. The tool visualizes the resulting decision tree to analyze splits it makes in the data, the predicted classes in comparison with ground truth, and–as a flow diagram—the way in which items of different classes are differentiated as they move through the decision tree. While designed for decision trees, most components of Boababview can also be applied to the random forest model we use in this work. The python package [37] for visualizing the flow of items through the decision tree can also visualize individual trees of a random forest. However, our focus is on exploratory analysis of cohorts rather than model building, which is why we do not rely on visualizations for in-depth model analysis. Infuse [20] is a visual analytics system for selecting attributes for classification models. The large overview visualization ranks attributes by various means of attribute selection algorithms to separate informative from non-informative attributes for the classification process. In Kokiri, we do not use attribute selection algorithms but rank attributes by an importance measure computed from the decisions made by the random forest model. The ranking thus directly reflects the classification model.

Most closely related to our work is the approach by Rauber et al. [30], who built on work by Brandoli et al. [6]: In the first step, lowdimensional representations of the data are shown to visually predict class separability and, presumably, classification performance. To improve separability, attribute subsets can be selected based on their importance in a random forest model. We, in contrast, first create the model and then create a low-dimensional representation of its decisions so that the visualization directly reflects the model. Good separation by the model results in good separation of the scatterplot, and items for which the model’s prediction is uncertain are identifiable as outliers. Additionally, we give a preview of the class distribution for each attribute when ranking them.

## 3 Kokiri

Kokiri compares and characterizes subsets of tabular data, here cohorts.The design and development process included regular feedback sessions with two experts working in a drug discovery team at a pharmaceutical company. The workflow within Kokiri, as shown in Fig. 2, consists of three steps that correspond to its three views: (i) the *Overlap View* to see which cohorts share items; (ii) the *Comparison View* to compare cohorts and obtain a ranking of the most important attributes and an overview of the classification performance; and (iii) the *Characterization View* to examine cohort homogeneity and hard-to-classify items. The individual cohorts are colored consistently across all visualizations using a categorical color schema (D3’s *Set 3* [5], see Fig. 1).

**Figure 1.**
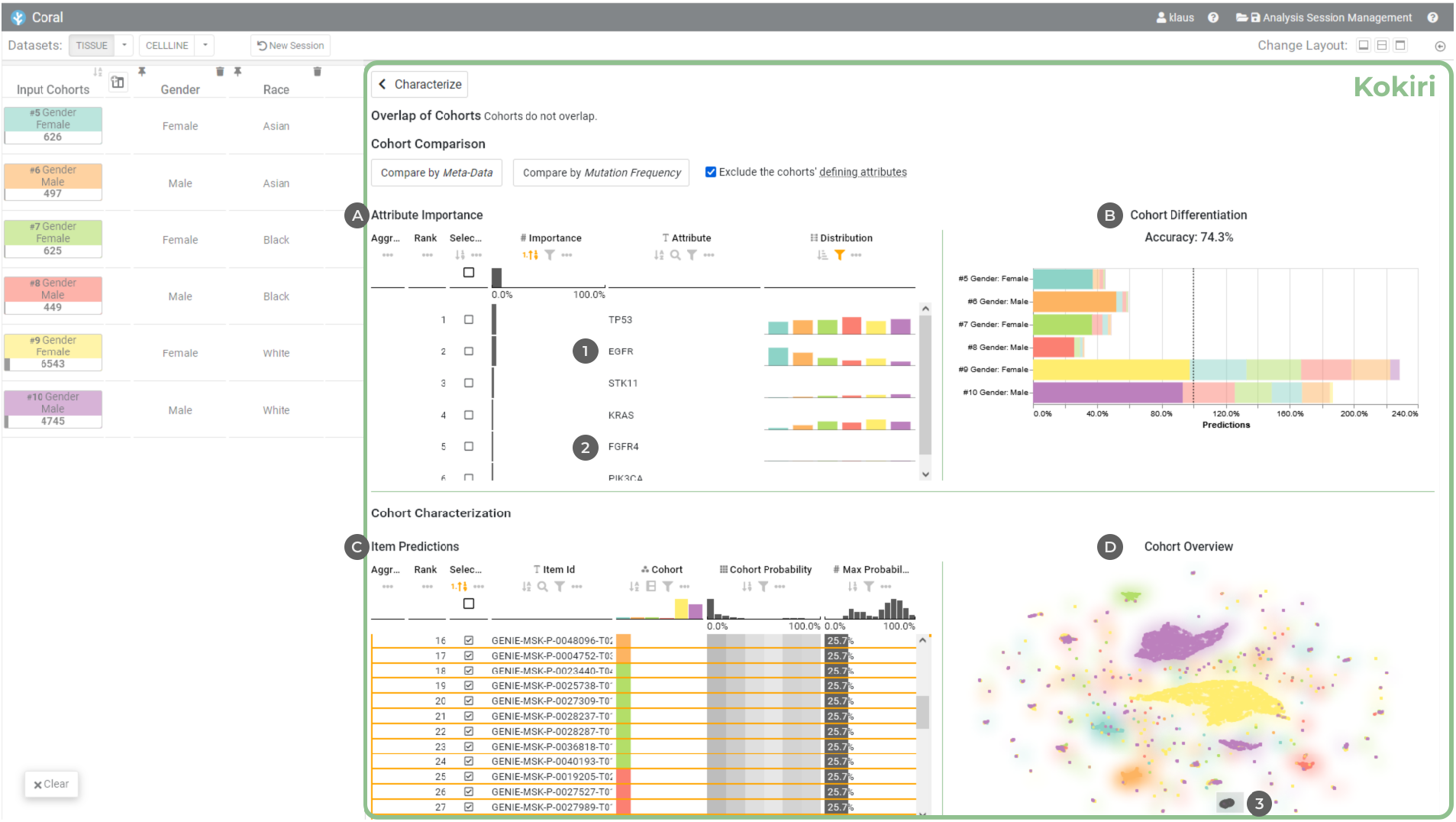
Kokiri integrated into Coral [2] comparing lung cancer patient cohorts of different races and genders. The ranked list **A** shows the most important attributes for differentiating the cohorts. The overall separability is shown with stacked bar charts **B**. A scatterplot **D** gives an overview of the predicted cohort affiliations of the patients, and a second ranked list C displays all items, the cohorts they belong to, and the probabilities to belong to any of the cohorts based on the data.

**Figure 2:**
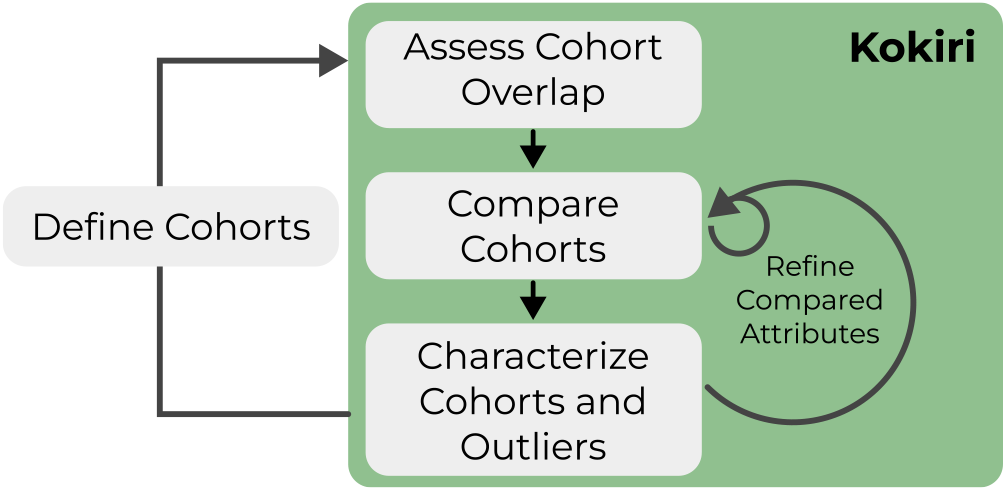
The workflow in Kokiri starts with assessing the overlap between cohorts. The cohorts are then compared to find the most important differentiating attributes. In the last step, cohort homogeneity and outliers are characterized. Based on their findings, users can compare cohorts on a different set of attributes or refine cohorts outside of Kokiri.

To compare and characterize the cohorts, we train a random forest classifier [7] that aims to differentiate them based on the data provided. A random forest is an ensemble of decision trees that improves generalization by introducing randomness to the training process: (a) each decision tree is trained on a bootstrapped sample of the data; that is, each tree sees a different subset of the data; and (b) only a random subset of attributes is considered at each split. To which cohort an item belongs is determined by averaging the decisions of the forest’s trees. Random forests have the advantages that they are interpretable, work well for biomedical data, and can capture complex interactions in the data [9,16,27].

### Overlap View

The Overlap View shows the pairwise overlap between cohorts that share items. Each cohort is represented by a bar, and bars grow in opposite directions. The length of each bar corresponds to the number of items in the cohort relative to the number of items in both cohorts. For example, the overlap of a cohort 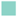 with 160 items and a cohort 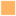 with 320 items that share 80 items (20% of the total items) is visualized as: 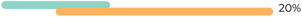 The visualization thus shows the relative overlap and differences in size, similar to a collapsed representation of an Euler diagram and the visualization proposed by Borland et al. [4]. In the context of Kokiri, we have found this visualization better suited than the alternatives tested—including UpSet [22], Jaccard Similarity [18], and absolute counts of intersecting items—due to its compactness and representation of relative cohort sizes.

We highlight the overlap of cohorts at the top of Kokiri’s interface because overlapping cohorts are harder to distinguish, items may be part of multiple cohorts, and the random forest model may assign items to multiple cohorts. As a result, the overlaps also affect the Comparison and Characterization Views of Kokiri.

### Comparison View

We iteratively train a random forest classifier, increasing its number of estimators in steps of 25 up to a total of 500 to give early feedback on the performance and the most important attributes. This allows users to stop the training early, giving them a chance to incorporate their domain knowledge by excluding already known differences or focusing on a subset of attributes for the next comparison. Human and machine pattern recognition capabilities can thus be combined, and users gain a deeper understanding of the resulting classification model and can also actively improve it [3,12,23]. Attributes important to the classification are determined by their Gini importance, which is an attribute importance measure that describes the quality of a split on an attribute: The better the split, the higher the Gini importance. The values of all splits from the same attribute are summed per tree and averaged over the whole forest. As the Gini importance is used to select the best attribute for splitting the data, it is already computed while the forest is built [27].

The *Comparison View* gives insights into how well the cohorts are separable by the data and which attributes are most able to differentiate them. It consists of two components that are displayed side-by-side: (i) a ranking of the attributes based on importance; and (ii) an overview of the classifier’s performance.

We use a tabular visualization [15] in which the attributes are ranked by their importance for distinguishing the cohorts (see Fig. 1A). Numerical attributes are represented by a row in the table. Categorical attributes have one row per category, but can be grouped together by the user. By having one row per category, we can show with a bar chart how many items of each cohort fall within this category. This distribution is shown in a separate column so that users can understand why an attribute is important to the classifier. For numerical data, we visualize the distribution of the cohorts using density plots.

The performance overview shows the mean accuracy score and the composition of the cohorts as predicted by the classifier (see Fig. 1 B). We visualize the predicted composition with a stacked bar chart. Each cohort is represented by a bar, and colored segments represent the items the classifier assigned to the cohort. The segments are ordered by their size and segments of misclassified items have lower opacity.

### Characterization View

To characterize cohorts and outliers, we rely on the classification model trained in the previous comparison step. To predict the cohort an item belongs to, each tree of the random forest casts a vote, and the majority vote decides the class. The more the votes differ, the less confident the classifier is about assigning an item to a cohort, and the lower the probability that an item belongs to a cohort. For *N* items and *k* cohorts, the probabilities form a vector of size *N* × *k*. Using this vector, we can characterize items that are (a) clearly assignable to a cohort, (b) assignable to a cohort but with low probability because the data deviate, (c) equally assignable to several cohorts, and (d) assignable to no cohort (i.e., outliers).

We visualize these probabilities following an overview+detail approach with: (i) a ranking of items based on the prediction probability to quickly identify those that the classifier was most uncertain about, and (ii) an embedding scatterplot that gives an overview of the cohort’s homogeneity, potential sub-cohorts, and outliers.

As for the attribute ranking, we use a tabular visualization to present the item ranking. Initially, the items are ranked in ascending order according to the maximum in the prediction probability vector. The table shows the individual items, the cohorts to which they belong, and to what degree this assignment was predicted (see Fig. 1 c).

For the embedding scatterplot, the prediction probability vector described above is reduced to two dimensions by means of supervised UMAP [24]. Supervised UMAP preserves the structural relationships, but reduces the overlap of classes within clusters. This allows misclassified items to be identified more easily because overplotting by other classes is reduced. Beneath the scatterplot marks, we display a heatmap that shows the cohort with the highest prediction probability in the respective area with the cohort’s color (see Fig. 1 D). In the heatmap, opacity increases with increasing classifier confidence, that is, when the probability of the predicted cohort differs most from those of the other cohorts. Thus, items that can be confidently assigned to a cohort are in an area with that cohort’s color, while items that can be assigned less confidently have little to no background. Users can select items in the table or the scatterplot. The selections are synchronized such that outliers and subgroups can be explored from both visualizations.

## 4 Implementation

Kokiri uses a client-server architecture. The server-side is written in Python, reads the data from a DuckDB [28], and trains a random forest classifier using scikit-learn [26]. The client-side is a web component written in TypeScript. We use the LineUp technique [15] for the rankings of attributes and items and Vega [31] to visualize the performance overview and the embedding of prediction probabilities in a scatterplot. Kokiri is open-source and available on GitHub: https://github.com/jku-vds-lab/kokiri/.

We have integrated Kokiri into Coral [2] as a *Characterize* operation to compare cohorts, either by the metadata or the genomics data, including mutation status. The prototype integration of Kokiri into Coral is available at https://kokiri.jku-vds-lab.at/.

## 5 Use Case

In this use case, we used the prototype integration of Kokiri in Coral to analyze lung cancer patient cohorts of different races and genders. The workflow and interactions are demonstrated in the supplementary video. A recent study [25] has shown differences in *KRAS G12C* mutation frequency between these cohorts. The discovery was based on the AACR Project GENIE [1] data, and we were able to reproduce it in Coral [2]. With Kokiri, we want to investigate whether there are any further mutational differences of interest. The dataset includes 112935 patients for whom metadata such as age, gender, and tumor type were recorded, as well as information on 3252 genetic mutations. For the comparison in Kokiri, we consider mutations that result in an amino acid change in any of 719 genes that were identified as tumor-relevant in the Cancer Gene Census [34].

We start the analysis from a previous session in Coral^1^ in which the relevant cohorts have already been created. Coral’s cohort evolution view shows that the dataset was filtered to non-small cell lung cancer tumor patients and subsequently split into cohorts of Asian, Black, and White patients and further into Female and Male patients. We select Female and Male cohorts for the three races and open Kokiri in Coral. As the cohorts are created by split operations, the *Overlap View* shows only a short note that the cohorts do not overlap. In the *Comparison View*, the cohorts can be compared by their metadata or genetic mutations. Comparison using metadata excludes by default all attributes used to create the cohorts. We compare the cohorts by genetic mutations, which starts training of the classifier and shows intermediate results after the first 25 decision trees of the random forest have been trained.

The initial results show that the genes *TP53, EGFR, STK11, KRAS*, and *FGFR4* differ most between these cohorts. In the overview of the classifier’s performance, we can also see that for the White Female and Male cohorts—the two largest cohorts—most items are correctly assigned. However, large proportions of patients from the other cohorts are also assigned to these two cohorts (see Fig. 1 B). The distribution column of the attribute ranking shows how many patients have a mutation in these genes. As expected, the mutation frequency of *TP53* is high in all cohorts, and it is also the most mutated gene in the whole dataset. We can also see that Male cohorts have more *TP53* mutations than the Female cohorts of the same race. The opposite is observed in the distribution of EGFR mutations, where the mutation is more prevalent in Female than in Male patients of the same race. A much clearer difference, however, is seen in the distribution of *EGFR* mutations between the two Asian cohorts and the others (see 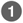 in Fig. 1). Differences in *EGFR* mutation frequency between Asian Male and Asian Female lung cancer cohorts and between Asian and White cohorts have also been described in a study by Shi et al. [32]. The frequency of *KRAS* gene mutations mirrors the distribution of *KRAS G12C* mutations described by Nassar et al. [25]. The distribution of *FGFR4* mutations, for which the bars are hardly visible (see Fig. 1 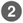), is also noteworthy. We investigate the distribution of *FGFR4* mutations further and see that there is indeed little difference in the number of mutations between cohorts (see Fig. 3). However, *FGFR4* has been sequenced much less frequently for the two Black cohorts. Presumably, there would be no differences between the cohorts if the same amount of data were available. Nevertheless, the difference is noteworthy because it raises questions about the data.

**Figure 3:**
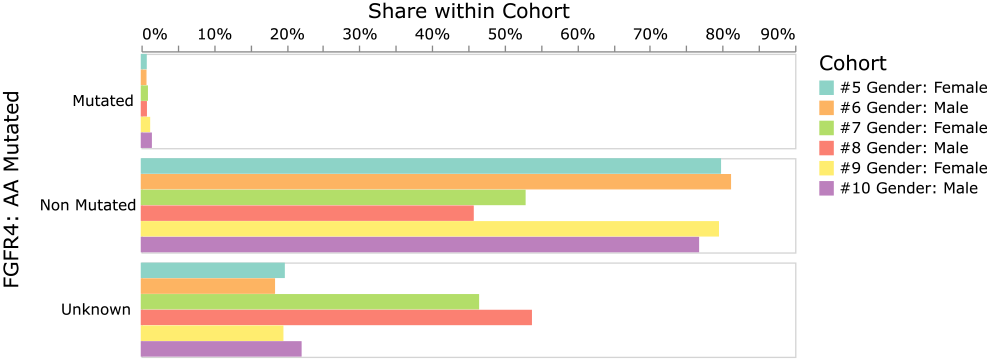
Differences in *FGFR4* mutation in Coral’s View operation. Black Female 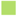 and Male 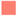 lung cancer patients have had *FGFR4* sequenced much less frequently than the other cohorts.

To further analyze the trained classifier and the confusion of Asian and Black cohorts with the two White cohorts, we proceed with the *Characterization View*. The embedding scatterplot shows not only a distinct cluster for each of the six cohorts, but also many smaller clusters as well as single items. By hovering over one of the smaller clusters (see 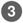 in Fig. 1), we see that the probability of being assigned to the White Female cohort was just 26%. We select the items in this cluster, which also selects them in the item ranking, where we can also see probabilities for all other cohorts. Even though the items in the selected cluster originate from different cohorts, the prediction probabilities are very similar, which suggests that their data is also similar. Ranking the items again by maximum probability, we see further items that the classifier was uncertain about. This could be a point in the workflow where users leave Kokiri and define new cohorts based on the findings (see Fig. 2).

To examine whether there are other differences in the data collected besides *FGFR4*, we select the overall non-small cell lung cancer cohort and the Black patient cohort. Kokiri’s overlap visualization shows that the Black cohort 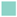 is a subset of the non-small cell lung cancer cohort 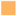, and that it is also much smaller:

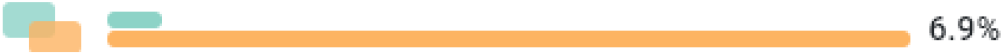

By comparing the cohorts by mutation frequency, we can now see that *FGFR4* is the most important attribute to classify Black patients. *BRCA1* is also among the most important genes, and the bars in the distribution column are hardly visible, as seen for *FGFR4* in Fig. 1 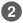. We find that this gene has also been sequenced less frequently for Black patients: For 50% the mutation status is unknown, compared to 25% in the entire non-small cell lung cancer cohort. One reason may be the huge differences in how comprehensively genomics data are collected at the different facilities at which the patients are treated.

## 6 Conclusion

In this paper, we have presented Kokiri, a visual analytics approach to comparing patient cohorts by their high-dimensional data, exploring the driving differences between cohorts, and characterizing the homogeneity and outliers of a cohort. Kokiri can compare multiple cohorts at once and captures combinatorial effects in the data by training a random forest classifier to distinguish between the cohorts.

Kokiri can be applied conceptually and technically to tabular data from other fields, for example, to explain differences between clusters in low-dimensional embeddings. We plan to integrate Kokiri into the Projection Space Explorer [17] for this purpose.

We also plan to enhance the robustness of the classifier and analyze more specific implementations of random forest classifiers. We believe that differences in missing data—as shown in the use case above—can be informative, but recognize the need to give users the ability to omit their consideration. The heterogeneity we saw in the use case, presumably due to different origins of the data, is also a concern when comparing cohorts of multiple datasets. Recent work [29, 39] has investigated strategies for adapting random forests to better handle heterogeneous data. Strobl et al. [36] pointed out a bias in the random forest attribute importances and suggested an implementation that provides unbiased results.

## Acknowledgments

The authors acknowledge the American Association for Cancer Research and its financial and material support in the development of the AACR Project GENIE registry, and members of the consortium for their commitment to open data. Interpretations are the responsibility of study authors. This work was supported in part by the Boehringer Ingelheim Regional Center Vienna, the State of Upper Austria, and the Austrian Federal Ministry of Education, Science and Research via the LIT – Linz Institute of Technology (LIT-2019-7-SEE-117), the State of Upper Austria (Human-Interpretable Machine Learning), and the Austrian Science Fund (FWF DFH 23-N).

## Footnotes

1 http://vistories.org/kokiri-use-case

